# *Streptococcus pyogenes* forms serotype and local environment-dependent inter-species protein complexes

**DOI:** 10.1101/2021.02.09.430411

**Authors:** Sounak Chowdhury, Hamed Khakzad, Gizem Ertürk Bergdahl, Rolf Lood, Simon Ekstrom, Dirk Linke, Lars Malmström, Lotta Happonen, Johan Malmström

## Abstract

*Streptococcus pyogenes* is known to cause both mucosal and systemic infections in humans. In this study, we used a combination of quantitative and structural mass spectrometry techniques to determine the composition and structure of the interaction network formed between human plasma proteins and the surface of different *S. pyogenes* serotypes. Quantitative network analysis revealed that *S. pyogenes* form serotype-specific interaction networks that are highly dependent on the domain arrangement of the surface-attached M protein. Subsequent structural mass spectrometry analysis and computational modelling on one of the M proteins, M28 revealed that the network structure changes across different host microenvironments. We report that M28 binds secretory IgA via two separate binding sites with high affinity in saliva. During vascular leakage mimicked by increasing plasma concentrations in saliva, the binding of secretory IgA was replaced by binding of monomeric IgA and C4BP. This indicates that an upsurge of C4BP in the local microenvironment due to damage of the mucosal membrane drives binding of C4BP and monomeric IgA to M28. The results suggest that *S. pyogenes* has evolved to form microenvironment-dependent host-pathogen protein complexes to combat the human immune surveillance during both mucosal and systemic infections.

## Introduction

Bacterial pathogens have evolved to express a multitude of virulence factors on their surface to establish versatile host-pathogen protein-protein interactions (HP-PPI)^1^. These interactions range from binary interactions between two proteins to the formation of multimeric interspecies protein complexes that enable bacterial pathogens to hijack and re-wire molecular host systems to circumvent host immune defenses. One prominent example is *Streptococcus pyogenes*, a gram-positive and beta-hemolytic bacterium. This bacterium causes diverse clinical manifestations such as mild and local infections like tonsillitis, impetigo and erysipelas as well as life-threatening systemic diseases like sepsis, meningitis and necrotizing fasciitis^2^. Globally, 700 million people suffer from *S. pyogenes* infections every year leading to an estimated 160,000 deaths^3^, thus making *S. pyogenes* one of the most widespread bacterial pathogens in the human population. *S. pyogenes* abundantly produce a prominent surface antigen, the M protein, known to enable bacterial invasion into human cells, prevent phagocytosis ^4,5^ and promote survival in infected tissues^6,7^. These M proteins are dimeric α-helically coiled-coil proteins covalently attached to the *S. pyogenes* cell wall and extending approximately 500 Å into the extra-bacterial space to form a dense fibrillary coat on the bacterial surface^8^. The M proteins consist several protein domains, some of which are repeat regions (**Fig 1A**). The N-terminal 50 amino acid residues constitute the hypervariable region (HVR)^9,10^. Sequence variation within the HVR is used to classify the M protein and till date >220 distinct *S. pyogenes* serotypes have been reported^8^. The HVR is followed by a stretch of 100-150 amino acids that forms the semi-variable domain of the M proteins and encompasses the A domain and the B repeats. The subsequent C repeats and the D domain form the conserved C-terminal part of the M proteins. The M proteins are classified into different *emm*-patterns *e.g.* A-C, D and E based on the arrangement of the A, B, C and D domains. The *emm* pattern A-C represent long M proteins with A, B, C and D domains, the *emm* pattern D includes M proteins with B, C and D domains, while the *emm* pattern E only includes the C and D domains^11,12^ (**Fig 1A**). It has been reported that the *emm* pattern A-C mainly includes *S. pyogenes* strains associated with throat infections, the *emm* pattern D includes *S. pyogenes* strains responsible for skin infections, while the E pattern includes generalist *S. pyogenes* strains typically infecting both sites^12^, indicating that the M protein domain composition correlates with host tissue tropisms. Furthermore, comparative sequence analysis of the M proteins enables classification of the M proteins into clades. Clade X includes the E pattern and clade Y the A-C pattern, while pattern D seem to fall into both clades X and Y^13^.

**Figure 1.**
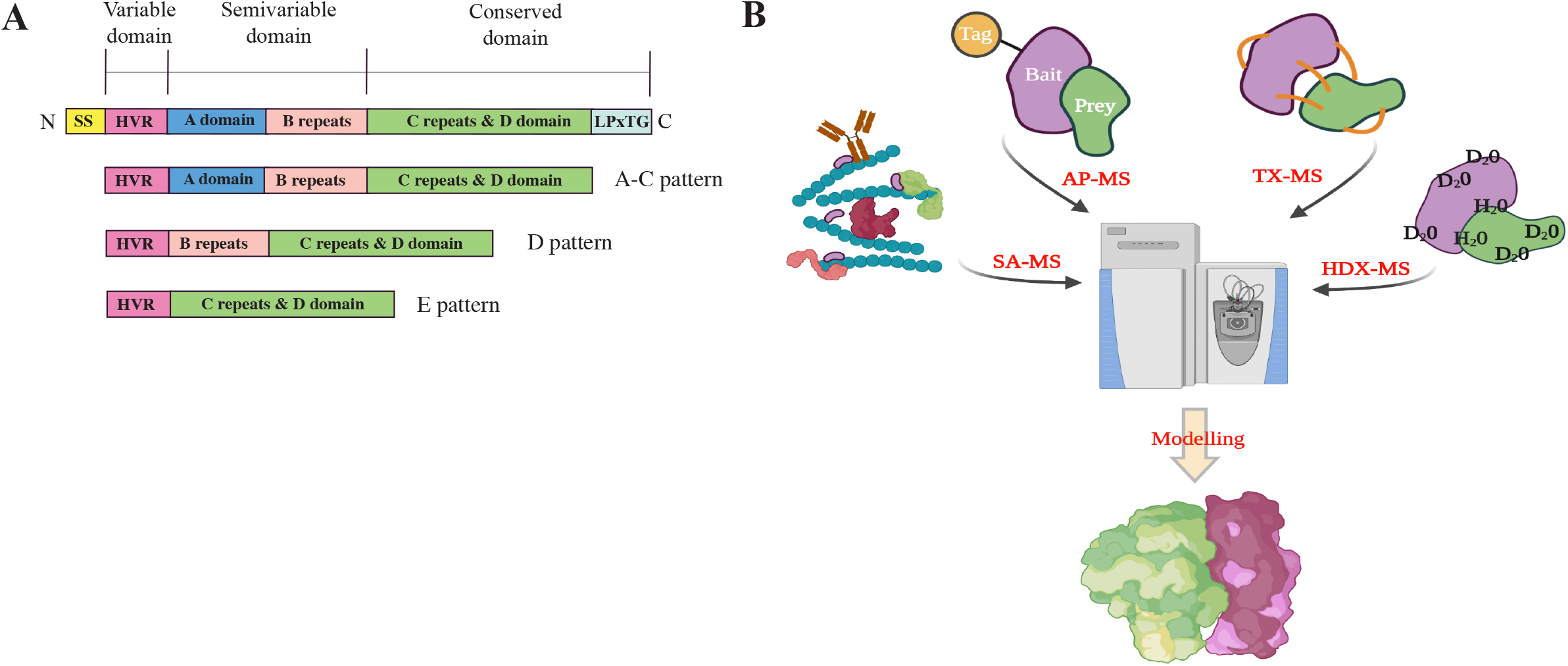
M protein (naïve and mature) structure and the experimental overview to identify *S. pyogenes*-human protein interactions. A) The arrangement of the different domains in M proteins. SS: signal sequence; HVR: hyper-variable domain, which is unique amongst different M proteins thereby giving rise to the numerous *S. pyogenes* serotypes; A domain and B repeats forming the semi-variable domain; C repeats and D domains including the LPXTG anchor sequence forming the conserved domain. Cleavage of the SS leads to a mature M protein and LPXTG helps the M protein anchor the bacterial surface. The S region of certain M proteins are not represented in this figure. M proteins are classified into A-C pattern harboring A domain, B and C repeats and the D domains, D pattern comprising the B-C repeats and the D domains and E type harboring only C repeats and the D domain. The M proteins are not drawn to scale. B) Schematic overview of the integrative approach used to characterize the protein network and complex around *S. pyogenes*. In SA-MS, pathogens are incubated with complex biological mixtures to capture proteins interacting with the bacterial surface which are then identified and quantified by MS. In AP-MS, recombinant bait proteins are expressed which are then made to capture prey proteins from complex biological mixtures followed by identification and quantification by MS. TX-MS is used to cross-link the protein partners to map the binding site while HDX-MS identifies the binding site based upon the exchange of H_2_O and D_2_O. Computational modelling is then used to generate protein interaction models based on the identified protein interaction sites.

The diverse domain arrangement and partially high sequence variability of the M proteins enables *S. pyogenes* to form protein interactions with various human proteins^14,15,16^. A recent chemical cross-linking mass spectrometry and structural modelling study showed that the M1 protein of the *emm* pattern A-C is capable of forming a large 1.8 MDa interspecies protein complex with up to 10 different human proteins^17^. The model by Hauri *et al.* show that the interacting human proteins are precisely placed along the α-helically coiled-coil structure of the M1 protein. In this way, the M protein can form a highly organized human plasma protein interaction network on the bacterial surface consisting of both human-human and *S. pyogenes*-human protein interactions^18^. One example is the binding of fibrinogen to the B repeats of the M protein^10,14,19^, where fibrinogen in turn mediates binding to factor thirteen (F13). Fibrinogen binding to the M protein prevents deposition of opsonizing antibodies to inhibit phagocytosis^20–22^. Several copies of human serum albumin (HSA) have been proposed to bind the C repeats of the M proteins to facilitate the uptake of fatty acids and promote growth during stationary phase ^14,20,23,24^. Additionally, certain *S. pyogenes* serotypes can bind immunoglobulin G (IgG). The orientation of IgG binding *i.e.*, whether Fab- or Fc-mediated is governed by the concentration of IgGs in the host niche^15,25^. The binding to the IgG-Fc is mediated by the S region found in some M proteins and located between the B and C repeats and the HVR region^14,26^. Other M proteins such as M4 and M22 of the *emm* pattern E, have been shown to bind immunoglobulin A (IgA)^15,27^. This binding occurs between the N-terminus of M proteins ^28,29^ and the inter-domain region of IgA-Fc, which is also the known binding site of human IgA receptor CD89^30^. Binding of M proteins to IgA-Fc is believed to block the binding of IgA to CD89 and thus prevent IgA-effector functions, inhibit phagocytosis and promoting bacterial virulence^30,31^. In addition, many different *emm* pattern M proteins bind complement system C4b-binding protein (C4BP) to the N-terminal HVR domain^32–36^. C4BP bound to the M protein sequesters C4b from plasma, and acts as a co-factor for the degradation of C4b by complement factor I^33,37,38^, thereby inhibiting the classical complement pathway and phagocytosis of the bacterium^31^.

Collectively, these previous studies indicate that the domain arrangement of different M proteins impacts the HP-PPI networks that are formed around the streptococcal surface. However, the large variability between different M proteins, the difficulty to pin-point exact binding interfaces and the formation of human-human protein interactions at the streptococcal surface makes it challenging to determine the structure and composition of such interspecies protein networks. More detailed understanding of how the domain arrangement determines the composition of M protein-centered interspecies proteins complexes could help explain differences in tissue tropism observed between different *emm* types. Here we applied quantitative and structural mass spectrometry techniques in an unbiased fashion to show that different serotypes form highly distinct *emm* pattern-specific HP-PPI networks. These interaction networks depend to a large extent on the type of M proteins expressed by a given strain. Furthermore, by in-depth structural mass spectrometry and structural modelling analysis, we demonstrate that the M proteins are capable of altering the composition of the protein complexes depending on the local microenvironments to enable critical immune evasion strategies in different ecological niches.

## Methods

### Cloning, expression and purification of recombinant proteins

The M proteins were cloned, expressed and purified at the Lund Protein Production Platform (LP3; Lund, Sweden) and at the University of Oslo (Norway). The recombinant M proteins used in this study lacked the signal peptide and the LPXTG cell wall-anchoring motif. The open reading frames corresponding to the mature M proteins - M1 (Uniprot ID: Q99XV0; aa 42-448, gene name: *emm1*), M3 (Uniprot ID: W0T370; aa 42-545, gene name: *emm3*), M5 (Uniprot ID: P02977; aa 43-456, gene name: *emm5*), M28 (Uniprot ID: W0T1Y4; aa 42-358, gene name: *emm28*), M49 (Uniprot ID: P16947; aa 42-354, gene name: *emm49*) and M89 (Uniprot ID: W0T3V8; aa 42-360, gene name: *emm89*) were cloned into a pET26b(+)-derived vector carrying 6xHis-HA-StrepII-TEV (histidine-hemagglutinin-StrepII-tobacco etch virus protease recognition site)tag. The protein sequences are provided in supplementary table 1 (**ST1**). These proteins were expressed in *E. coli* Tuner (DE3) induced with 1mM isopropyl β-D-1-thiogalactopyranoside (IPTG) at an OD_600_ of 0.6 at 18°C, 120 rpm after 18 hours. The cells were harvested by centrifugation at 8000 g at 4°C for 20 min, and the pellets were re-suspended in 50 mM NaPO_4_, 300 mM NaCl, 20 mM imidazole, pH 8 (buffer A) supplemented with EDTA free complete protease inhibitor tablets (Roche). The cells were lysed using a French press at 18000 psi. The cell lysate was cleared by ultra-centrifugation at 45000 rpm for 60 min at 4°C with subsequent passing through a 0.45 μm syringe filter. The cell lysate was loaded onto a HisTrap HP column (GE Healthcare), followed by washing with 20 column volumes of buffer A and the bound proteins were subsequently eluted with a 0-100% gradient of buffer B (50 mM NaPO_4_, 300 mM NaCl, 500 mM imidazole, pH 8). The fractions containing the protein of interest were pooled and dialyzed against 1x phosphate-buffered saline (1xPBS; 10 mM phosphate buffer, 2.7 mM KCl, 137 mM NaCl, pH 7.3) and stored at −80°C until further use. The expression and purification of sfGFP have been described elsewhere^16^.

### Removal of the M28 affinity-tag by TEV-protease digestion

For SPR, targeted cross-linking mass spectrometry (TX-MS) and hydrogen-deuterium mass spectrometry (HDX-MS), M28 without the affinity-tag was used. For removal of the affinity-tag, the M28 protein was treated with TEV-protease at an enzyme: substrate mass ratio of 1:20. Dithiothreitol (DTT) was added to a final concentration of 1mM and the digestion mixture was transferred to a dialysis membrane (6-8000 molecular weight cut-off) and dialyzed against buffer A supplemented with 1 mm DTT at 16°C 18 hours. The mixture was passed through a HisTrap column (GE Healthcare) at room temperature with the same gradient and buffer as used earlier. Fractions containing the cleaved M28 protein were collected, and passed through a 0.2 μm syringe filter before loading on a 26/600 Superdex 200 pg column (GE Healthcare) run with 1xPBS (pH 7.4) at 2.5 ml/min at 6°C. Fractions with TEV-cleaved, purified M28 were pooled together and stored at −80°C until further use.

### Commercial proteins and human plasma and saliva

Pooled human plasma (LOT number 18944 and 27744) and pooled human saliva (catalog number IR100044P) was purchased from Innovative Research, USA. Pooled saliva was centrifuged at 1500 x g for 15min at 4°C followed by sterile filtration using 0.22 μm Steriflip filtration units (Millipore) and stored −20°C until further use. IgA from human serum (LOT number 0000085362) was purchased from Sigma-Aldrich, Germany. Purified human complement C4BP (catalog number A109, Lot number 4a) was obtained from Complement Technology, USA. Recombinant human IgA-Fc domain (catalog number PR00105) was purchased from Absolute Antibody, UK.

### Bacterial culture

*S. pyogenes* serotype M1 (SF370) was obtained from the American Type Culture Collection (ATCC; strain reference 700294), which was originally isolated from an infected wound. The other *S. pyogenes* serotypes M3, M5, M28, M49 and M89 used in this study were clinical isolates obtained from the blood of GAS infected patients at Lund university hospital and serotyped by clinical microbiology department of the hospital. These bacteria were grown on blood agar plates, and single colonies were isolated and grown in Todd-Hewitt (TH) broth supplemented with 0.6 % yeast extract at 37°C, in 5% CO_2_ 16 hours. Bacteria from the overnight culture were sub-cultured in TH broth with 0.6% yeast extract at 37°C, in 5% CO_2_ till mid-logarithmic phase (OD_600_ nm 0.4-0.5). The cells were harvested by centrifugation at 3500 g for 5 min. The pellets were washed in HEPES-buffer (50mM HEPES, 150 mM NaCl, pH 7.5) twice and re-centrifuged at 3500 g for 5 min. The washed cells were re-suspended in HEPES-buffer to a 1% solution. These cells were further used for SA-MS experiments.

### Bacterial surface adsorption of human plasma proteins

To capture human plasma proteins on *S. pyogenes* surface 400 μl of pooled normal human plasma was added to 100 μl of 1% bacterial solution in six biological replicates for each strain. The samples were vortexed briefly, and incubated at 37°C at 500 rpm for 30 min. Cells were harvested by centrifugation at 5000 g for 5 min, and washed three times with HEPES-buffer followed by centrifugations of 5000 g for 5 min, respectively. The cells were finally re-suspended in 100 μl HEPES-buffer. For limited proteolysis of surface-attached bacterial and human proteins, 2 μg of 0.5 μg/μl sequencing grade trypsin (Promega) was added, and the digestion was allowed to proceed at 37°C, 500 rpm for 60 min. The reaction was stopped on ice, and the supernatant collected by centrifugation at 1000 g for 15 min at 4°C. Any remaining bacteria in the supernatants were heat-killed at 85°C at 500 rpm for 5 min, prior to sample preparation for mass spectrometry.

### Affinity purification of human plasma and saliva proteins

For affinity purification (AP) reactions 20 μg of recombinant affinity-tagged M proteins was charged on Strep-Tactin Sepharose beads (IBA) equilibrated in 1x PBS. Affinity-tagged sfGFP was used as a negative control in all experiments. Pooled normal human plasma (100 μl) or saliva (200 μl) was then incubated with the protein-charged beads at 37 °C, 800 rpm, 1 h. Every 1ml of saliva was complemented with 10 μl protease inhibitor (Sigma). For saliva-plasma mixed environment experiments 100 μl saliva-plasma dilutions were made for 100% saliva, 1% plasma (99μl saliva + 1μl plasma), 10% plasma (90μl saliva + 10μl plasma) and 100% plasma and incubated with the protein-charged beads at 37 °C, 800 rpm, 1 h. The beads were washed with 10 ml ice-cold 1x PBS (for plasma) and 4 ml ice-cold 1xPBS (for saliva and saliva-plasma dilutions) at 4 °C, before eluting the proteins with 120 μl 5 mM biotin in 1xPBS at room temperature (RT). To remove biotin from the eluted protein mixture tri-chloro acetic acid (TCA) was added to a final concentration of 25% and incubated in −20C for 16hours. The protein mixture was centrifuged at 13000 rpm for 30 min at 4 °C. The pellets were washed two times in 500 μl and once in 200 μl ice-cold acetone by centrifuging at 13000 rpm 10 min at 4 °C. These pellets were then prepared for mass spectrometry.

### Crosslinking IgA-Fc and C4BP with M28

10 μg of C4BP and 10 μg of IgA-Fc was incubated separately with 10 μg of M28 in a final volume of 100 μl in 1X PBS for 30 minutes at 37°C and 800rpm. To cross-link IgA-Fc or C4BP to M28 heavy/light disuccinimidylsuberate crosslinker (DSS-H12/D12, Creative Molecules Inc, www.creativemolecules.com) resuspended in 100% dimethylformamide (DMF) was added to final concentrations of 0, 100, 250, 500, 1000 and 2000 μM. The cross-linking mixture was then incubated at 37°C, 800 rpm for 60 minutes. Before preparing the sample for MS analysis the reaction was quenched by adding ammonium bicarbonate to a final concentration of 50mM and incubating for 15 min at 37°C and 800 rpm.

### Sample preparation for mass spectrometry

To denature the proteins 8M urea-100 mM ammonium bicarbonate was added to the SA-MS AP-MS and cross-linked samples. The disulfide bonds were reduced with 500 mM TCEP at 37°C for 60 min, and then alkylated with 500 mM iodoacetamide in the dark at room temperature for 30 min. The samples were diluted with 100 mM ammonium bicarbonate for a final urea concentration <1.5 M, and then 0.5 μg/μl sequencing grade trypsin (Promega) was added for protein digestion at 37°C for 18h. Mass spectrometry samples for cross-linking reactions were prepared in a similar fashion as stated above with an additional step of digestion with 0.5 μg/μl lysl endopeptidase (Wako) at 37°C, 800 rpm for 2 hours after treatment with iodoacetamide followed by dilution with ammonium bicarbonate and trypsin digestion. The digestions were quenched with 10% formic acid to a final pH of 2-3. The peptides were purified in SOLAμ HRP 2mg/1ml 96 well plate (Thermo Scientific) according to manufacturer’s protocol. The eluted peptides were dried in a speedvac, and resuspended in 2% acetonitrile −0.1% formic acid with iRT peptides^39^ (retention time peptides-as internal reference), followed by 5 mins sonication and brief centrifugation before mass spectrometry.

### Liquid chromatography tandem mass spectrometry (LC-MS/MS)

The peptides were analyzed using data-dependent mass spectrometry analysis (DDA-MS) and data-independent mass spectrometry analysis (DIA-MS) on a Q Exactive HFX (Thermo Scientific) connected to an EASY-nLC 1200 (Thermo Scientific). The peptides were separated on a Thermo EASY-Spray column (Thermo Scientific 50cm column, column temperature 45 °C) operated at a maximum pressure of 800 bar. A linear gradient of 4% to 45% acetonitrile in aqueous 0.1% formic acid was run for 65 min for both DDA and DIA. For DDA analysis, one full MS scan (resolution 60,000 for a mass range of 390-1210 m/z) was followed by MS/MS scans (resolution 15,000) of the 15 most abundant ion signals. The precursor ions with 2 m/z isolation width were isolated and fragmented using higher-energy collisional-induced dissociation at a normalized collision energy of 30. The automatic gain control was set as 3e6 for full MS scan and 1e5 for MS/MS. For DIA, a full MS scan (resolution 60,000 for a mass range of 390-1210 m/z) was followed by 32 MS/MS full fragmentation scans (resolution 30,000) using an isolation window of 26 m/z (including 0.5 m/z overlap between the previous and next window). The precursor ions within each isolation window were fragmented using higher-energy collisional-induced dissociation at a normalized collision energy of 30. The automatic gain control was set to 3e6 for MS and 1e6 for MS/MS. The cross-linked peptides were analysed in DDA. For DDA analysis of cross-linked peptides one full MS scan (resolution 60,000 for a mass range of 350-1600 m/z) was followed by MS/MS scans (resolution 15,000) of the 15 most abundant ion signals within an isolation width of 2m/z.

### SA-MS and AP-MS data analysis

MS raw data were converted to gzipped and Numpressed mzML^40^ using the tool MSconvert from the ProteoWizard, v3.0.5930 suite^41^. All data analyses were stored and managed using openBIS^42^. SA-MS DDA data acquired spectra were analyzed using the search engine X! Tandem (2013.06.15.1-LabKey, Insilicos, ISB)^43^, OMSSA (version 2.1.8)^44^ and COMET (version 2014.02 rev.2) against an in-house compiled database containing the *Homo sapiens* and *S. pyogenes* serotype M1 reference proteomes (UniProt proteome IDs UP000005640 and UP000000750, respectively) complemented with common contaminants from other species, yielding a total of 22 155 protein entries and an equal amount of reverse decoy sequences. AP-MS DDA data was analyzed using the same search engines as above, against an in-house compiled database containing the *Homo sapiens* and *S. pyogenes* serotype M1 reference proteomes (UniProt proteome IDs UP000005640 and UP000000750, respectively) complemented with the all of the affinity-tagged M proteins and the sfGFP sequences as well as common contaminants from other species, yielding a total of 22162 protein entries and an equal amount of reverse decoy sequences. Fully tryptic digestion was used allowing two missed cleavages. Carbamidomethylation (C) was set to static and oxidation (M) to variable modifications, respectively. Mass tolerance for precursor ions was set to 0.2 Da, and for fragment ions to 0.02 Da. Identified peptides were processed and analyzed through the Trans-Proteomic Pipeline (TPP v4.7 POLAR VORTEX rev 0, Build 201403121010) using PeptideProphet^45^. The false discovery rate (FDR) was estimated with Mayu (version 1.07) and peptide spectrum matches (PSMs) were filtered with protein FDR set to 1% resulting in a peptide FDR < 1%.

The SA-MS and AP-MS DIA data were processed using the OpenSWATH pipeline^46^. For DIA data analysis, spectral libraries from the above DDA dataset were created in openBIS^42^ using SpectraST (version 5.0, TPP v4.8.0 PHILAE, build 201506301157-exported (Ubuntu-x86_64)) in TPP^47^. For DIA data analysis, raw data files were converted to mzXML using msconvert and analyzed using OpenSWATH (version 2.0.1 revision: c23217e). The RT extraction window was ±300 s, and *m*/*z* extraction was set at 0.05 Da tolerance. RT was then calibrated using iRT peptides. Peptide precursors were identified by OpenSWATH (2.0.1) and PyProphet (2.0.1) was used to control the false discovery rate of 1% at peptide precursor level and at 1% at protein level. Then TRIC^48^ was used to align the runs in the retention time dimension and reduce the identification error by decreasing the number of missing values in the quantification matrix. Further missing values were re-quantified by TRIC^48^. Resulting DIA data sets were analysed using Jupyter Notebooks (version 3.1.1). For the DIA data analysis proteins identified by more than 3 peptides and enriched with a log2 fold enrichment of >1 (two-fold) with an adjusted P-value <0.05 using the Student’s t-test were considered has true interactors. However, for the saliva-plasma dilution DIA data TRIC was not enabled. The intensities of the proteins were estimated by summing the intensities of the most intense three peptides for each protein relative to the total peptide intensities (without iRT) for that protein.

### Surface Plasmon Resonance (SPR) analysis of M protein

Binding experiments were performed on Biacore X100 (Cytiva Life Sciences, Uppsala, Sweden) with a control software version of v.2.0. All the assays were carried out on a Sensor CM5 gold chip (Cytiva Life Sciences, Uppsala, Sweden) at 25 °C. For the covalent immobilization of M1 and M28 molecules via amine groups on the gold surface Amine coupling kit (Cytiva Life Sciences, Uppsala, Sweden) containing EDC [1-Ethyl-3-(3-dimethylamino-propyl)carbodiimide] (75 mg/mL), NHS (N-hydroxysuccinimide) (11.5 mg/mL) and ethanolamine (1 M, pH: 8.5) was used.

The CM5 chip was docked into the instrument and the chip surface was activated following EDC/NHS protocol with PBS buffer as the running buffer before the immobilization procedure. The ligand (M1/M28) was injected for 7 min (flow rate: 10 μL/min) at a concentration of 0.01 mg/mL (in 10 mM acetate buffer, pH: 5.0) followed by an injection of 1.0 M ethanolamine for 7 min (flow rate: 10 μL/min) in order to deactivate excess reactive groups. Once the targeted immobilization level (≈ 2500 RU) was achieved no further immobilization was carried out. The flow channel_2 (active channel) was used for the ligand immobilization while the flow channel_1 (reference channel) was used as a reference to investigate non-specific binding. Response units were recorded from the subtracted channel (flow channel_2 – flow channel_1) which was then used to evaluate the results of analysis. For the IgA as analyte, concentration series including 0, 0.009375, 0.01875, 0.0375, 0.075, 0.15 and 0.3 μM were prepared. For the C4BP as analyte, concentration series between 0 and 96 nM were prepared. The analytes were injected into the active (Fc_2) and reference channels (Fc_1) at the same time. Triplicate injections were done for each concentration series. The association time was set to 120 s while the dissociation time was kept as 600 s. For the regeneration of the surface, 10 mM glycine-HCl (pH: 2.5) was used at a flow rate of 10 μL/min.

### Evaluation of SPR Analysis

For the evaluation of the analysis, the kinetic parameters were determined by Biacore Evaluation Software (v.2.0) in binding analysis based on curve-fitting algorithms which employs global fitting.

The data collected for each experiment was analysed according to 1-1 fitting model using the kinetic fitting programs that yields k_a_, k_d_ and K_D_ values and also fitting the data to heterogeneous binding model. Equilibrium binding analysis were performed by plotting the RU values measured in the plateau versus each concentration series.

First the binding was tested for the simplest 1-1 Langmuir binding model, which follows the equation:

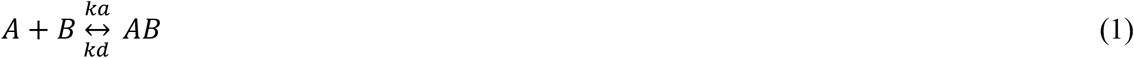

where A is the analyte, B is the ligand, AB is the complex. The k_a_ (rate of association, M^−1^s^−1^) is measured from the reaction in the forward direction while the k_d_ (dissociation rate, s^−1^) is measured from the reverse reaction.

The binding was also tested for heterogeneous ligand model where the same analyte binds independently to multiple ligands or to several binding sites on the same ligand. Heterogeneous ligand model follows the equation:

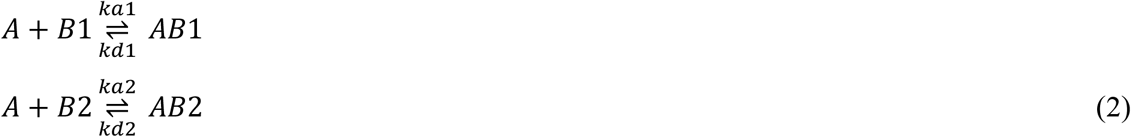

where A represents the analyte, B1 and B2 represent two different ligands or two different binding sites on the same ligand, respectively, AB1 and AB2 represent the first and second complexes formed after the binding of the analyte to the surface, ka_1_ and ka_2_ are the association rates of the first and second complexes while kd_1_ and kd_2_ represent the dissociation rates.

### TX-MS data analysis and computational modelling

The UniProt accession numbers used for the *S. pyogenes* M28 protein, human C4BPa, C4BPb, IGHA1, and IGHA2 were W0T1Y4, P04003, P20851, P01876, and P01877, respectively. The tertiary structure of the M28 protein was characterized using Rosetta comparative modeling (RosettaCM) protocol^49^ from Rosetta software suit ^50^ based on the previously generated full-length model of the M1 protein^26^ as the homologue structure. For IgA and C4BP, PDB ids 6LXW, and 5HYP have been used, respectively. To analyze the interactions of M28 with IgA and C4BP, the TX-MS protocol has been employed^17^ through which computational docking models were generated and filtered out using distance constraints derived from MS-DDA data. A final round of high-resolution modeling was performed on the top selected models to repack the sidechains using RosettaDock protocol^51^.

### HDX-MS sample preparation and data acquisition

HDX-MS was performed in two separate runs on M28 with IgA-Fc and M28 with C4BP. In each experimental run HDX-MS was first performed on pure untagged M28 (1 mg/mL). Then a mixture of M28 with the different ligands were prepared in PBS as described below and subjected to HDX.

M28: IgA(Fc)-1:1 molar ratio: Each sample consisted of 1 μl of M28 (75 pmol/μl) mixed with 1 μl PBS and 3 μl IgA(Fc) at concentration of 20 pmol/μl.

M28: IgA(Fc)-2:1 molar ratio: Each sample consisted of 2 μl of M28 (75 pmol/μl) mixed with 3 μl IgA(Fc) at concentration of 20 pmol/μl.

M28 (pure): In this run each sample consisted of 1 μl of M28 (75 pmol/μl) mixed with 4 μl PBS.

M28:C4BP: In this run each interaction sample consisted of 2 μl of M28 (75 pmol/μl) mixed with 5 μl C4PB (1-2 pmol/μl).

M28 (pure): each sample consisted of 2 μl of M28 (75 pmol/μl) mixed with 5 μl PBS.

The HDX-MS analysis was performed using automated sample preparation on a LEAP H/D-X PAL™ platform interfaced to an LC-MS system, comprising an Ultimate 3000 micro-LC coupled to an Orbitrap Q Exactive Plus MS. Samples of M28 with and without ligand were diluted with 25 μl 10 mM PBS pH 7.4 (for t = 0 samples) or with 25 μl HDX labelling buffer comprising dPBS same composition prepared in D_2_O, and pH adjusted to pH_(read)_ 7.0 with DCl diluted in D_2_O. The HDX reactions were carried out for t = 30, 300, 3000 at 20°C. The labelling was quenched by dilution of the labelled sample with 30 μl of 1% TFA, 0.4 M TCEP, 4 M Urea, at 1°C, 50 μl of the quenched sample was directly injected and subjected to online pepsin digestion at 4 °C on an in-house packed (POROS AL 20 μm immobilized pepsin) pepsin column, 2.1 x 30 mm. The online digestion and trapping was performed for 4 minutes using a flow rate of 50 μL/min with a running buffer of 0.1 % formic acid, pH 2.5. The peptides generated by pepsin digestion were subjected to on-line SPE on a PepMap300 C18 trap column (1 mm x 15mm) and washed with 0.1% FA for 60s. Thereafter, the trap column was switched in-line with a reversed-phase analytical column, Hypersil GOLD, particle size 1.9 μm, 1 x 50 mm, and separation was performed at 1°C using a gradient of 5-50 % B over 8 minutes and then from 50 to 90% B for 5 minutes, the mobile phases were 0.1 % formic acid (A) and 95 % acetonitrile with 0.1 % formic acid (B). Following the separation, the trap and column were equilibrated at 5% organic content, until the next injection. The needle port and sample loop were cleaned three times after each injection with mobile phase 5%MeOH and 0.1%FA, followed by 90% MeOH and 0.1%FA and a final wash of 5%MeOH and 0.1%FA. After each sample and blank injection, the Pepsin column was washed by injecting 90 μl of pepsin wash solution 1% FA /4 M urea /5% MeOH. In order to minimize carry-over a full blank was run between each sample injection. Separated peptides were analysed on a Q Exactive Plus MS, equipped with a HESI source operated at a capillary temperature of 250 °C. For undeuterated samples (t = 0s) 1 injection was acquired using data dependent MS/MS HCD for identification of generated peptides. For HDX analysis (all labelled samples and one t= 0s) MS full scan spectra at a setting of 70K resolution, automatic gain control 3e6, Max IT 200ms and scan range 300-2000 Da were collected.

### HDX-MS Data analysis

PEAKS Studio X. (Bioinformatics Solutions Inc., Waterloo, Canada) was used for peptide identification after pepsin digestion of undeuterated samples (i.e. timepoint 0 s.). The search was done on a FASTA file comprising the only the sequences of the analysed proteins, search criteria was a mass error tolerance of 15 ppm and a fragment mass error tolerance of 0.05 Da and allowing for fully unspecific cleavage by pepsin.

Peptides identified by PEAKS with a peptide score value of log P > 25 and no modifications were used to generate peptide lists containing peptide sequence, charge state and retention time for the HDX analysis. HDX data analysis and visualization was performed using HDExaminer, version 3.01 (Sierra Analytics Inc., Modesto, US). Due to the comparative nature of the measurements, the deuterium incorporation levels for the peptic peptides were derived from the observed mass difference between the deuterated and non-deuterated peptides without back-exchange correction using a fully deuterated sample. HDX data was normalized to 100% D_2_O content with an estimated average deuterium recovery of 75%. The peptide deuteration was determined from the average of all high and medium confidence results, with the two first residues of each peptide set to be unable to retain deuteration. The allowed retention time window was set to ± 0.5 minutes. Heatmaps settings were uncoloured proline, heavy smoothing and the difference heatmaps were drawn using the residual plot as significance criterion (±1 Da). The spectra for all timepoints were manually inspected; low scoring peptides, e.g. obvious outliers and peptides were retention time correction could not be made consistent were removed.

## Results

### Human plasma protein interaction networks with *S. pyogenes* surface proteins

M proteins are long extended surface attached proteins with various combinations of A, B, C and D domains **(Fig. 1A**) that allow the M proteins to engage in numerous protein interactions simultaneously. While the protein interaction network formed around A-C patterns is relatively well described^16^, less is known about the protein interaction network organized around E pattern strains. Here, we combined quantitative and structural mass spectrometry techniques, to determine how the different M protein domains within E and A-C patterns influence the composition and structure of the human plasma-*S. pyogenes* interaction network (**Fig. 1B**). First, we selected three representative clinical isolates from *emm* pattern type A-C (M1, M3 and M5) and three from type E (M28, M49 and M89) and performed bacterial surface adsorption mass spectrometry analysis (SA-MS)^16^ as schematically shown in **Figure 1B**. In this analysis, the intact clinical isolates were incubated with pooled normal human plasma. Surface adhered proteins were enriched via centrifugation and quantified by data-independent mass spectrometry analysis (DIA-MS). The data was stringently filtered, resulting in the identification of in total 92 surface bound plasma proteins which were further grouped into six major protein families according to their functional roles, *i.e.*, apolipoproteins, cell adhesion and cytoskeleton proteins, coagulation, complement, immunoglobulins and other plasma proteins (**Fig. 2A)**. The quantitative data matrix across each strain show that there are marked differences in the HP-PPI networks formed on the streptococcal surface between the serotypes producing A-C or E type M proteins (**Fig. 2A).** The A-C pattern strains typically bind fibrinogen and components of the complement system, whereas the E pattern typically bind with various apolipoproteins, immunoglobulins and components from the complement and coagulation system such as C4BP and vitamin K-dependent protein S (PROS) (**Fig. 2A**). The analyzed strains were furthermore capable of forming distinct serotype-specific interaction networks, also within their respective A-C or E patterns. To objectively determine the major components of these networks, we used co-expression network analysis (**Fig. 2B**). This analysis revealed four highly connected protein clusters (highlighted using semi-transparent circles) of strongly correlating human proteins (blue lines, r^2^ > 0.9) that bind to one or two of the strains. The highest correlating proteins associated with M49 and M89 networks were several apolipoproteins such as APOH and APOC4 (**Fig. 2B)**. In contrast, M3, M5 and to some degree M1, bound fibrinogen whereas M28 predominately associates with several proteins such as C4BP, PROS, APOB and IgA (**Fig. 2B**). Interestingly, the network view also shows that there are several proteins that are negatively correlated (r^2^ < −0.6) as indicated by the red lines in **Figure 2B.** To further visualize these binding patterns, correlation plots were plotted for selected proteins pairs from each protein cluster (**Fig 2C**). As expected, strong correlations were observed between proteins belonging to the same protein cluster in **Figure 2B** such as APOH-APOC4, FIBB-FIBA, C4BPA-C4BPB, APOB-PROS, IGHA1-IGHA2, CO3-CO4A, C4BPA-PROS and IGHA1-C4BPA. In contrast, other proteins appear to bind significantly more to some strains such as fibrinogen and C4BP, where M1, M3 and M5 bind fibrinogen but not C4BP, while M28, M49 and M89 bind C4BP but not fibrinogen (**Fig. 2C**). Moreover, high levels of fibrinogen were related to low levels of several other proteins of the M28, M48 and M89 network such as IgA, PROS and components of apolipoproteins such as APOH (**Fig. 2C**). These results demonstrate that each strain can assemble strain-specific HP-PPI network and that there are substantial differences in the interaction networks between *emm* types.

**Figure 2.**
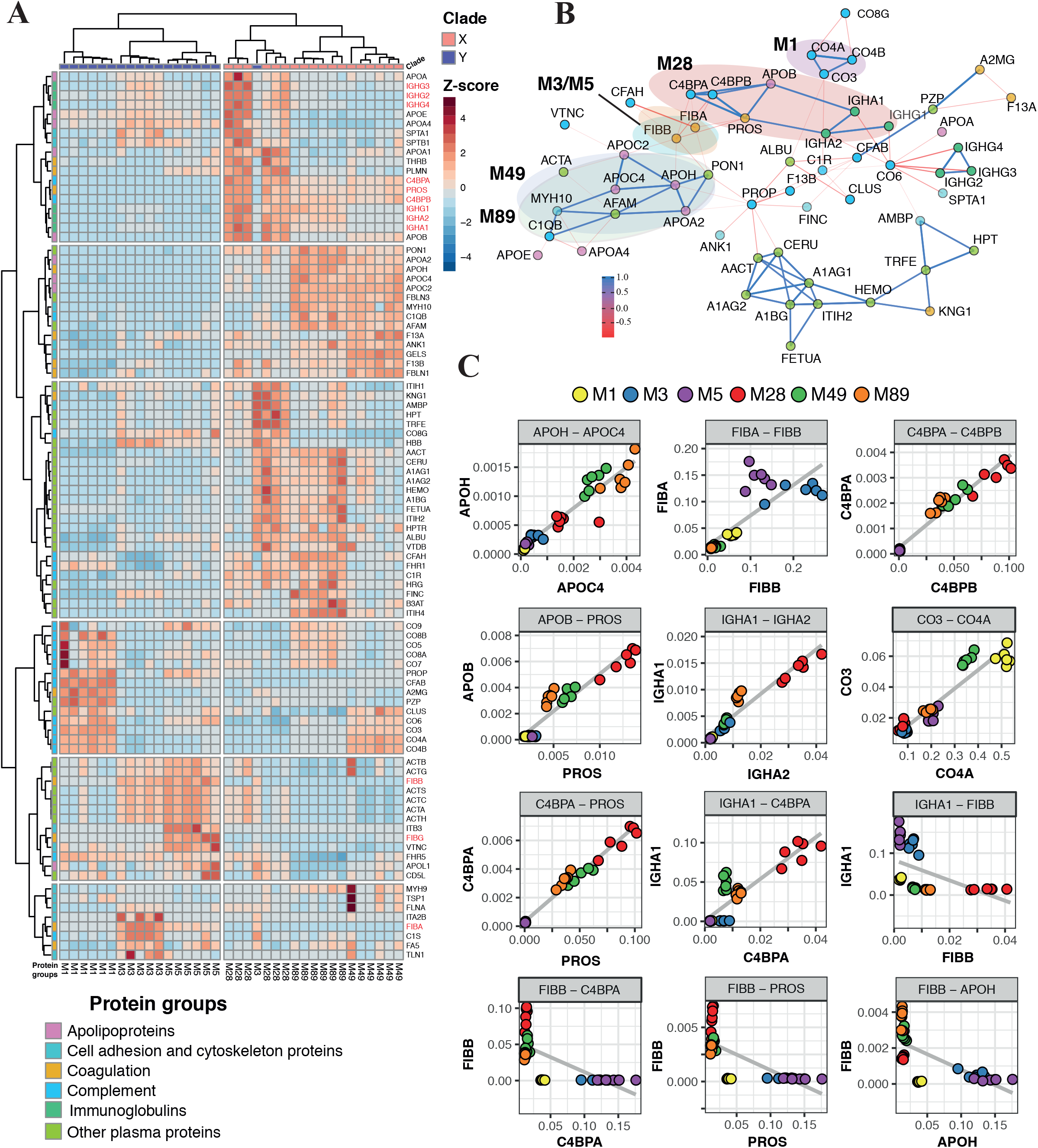
Human plasma proteins interacting with different *S. pyogenes* serotypes identified by SA-MS. A) Cluster analysis of 92 human plasma proteins interacting with six different *S. pyogenes* serotypes M1, M3, M5 (clade Y) and M28, M49 and M89 (clade X). SA-MS data is represented for n= 6 independent replicates for each strain. Human plasma proteins were categorized into six protein families, *i.e.*, apolipoproteins, cell adhesion and cytoskeleton proteins, coagulation, complement, immunoglobulins and other plasma proteins. The proteins colored in red are discussed in the text. B) Person correlation network analysis of human plasma proteins across six different serotypes. Each sphere represents a protein cluster. M3-M5 and M49-M89 share protein clusters, while M1 and M28 fall into individual clusters. Blue lines represent strongly correlating proteins (r^2^ > 0.9), while red line lines represent mutual exclusive proteins or negatively correlating proteins (r^2^ < −0.6). Protein dots are colored according to the protein family. C) Correlation plots for some representative proteins from each protein cluster across different strains. Strains are represented by dots with different colors.

### Human plasma protein interaction networks with *S. pyogenes* - M proteins

To understand to what degree differences in the domain arrangement of the M proteins (**Fig. 1A**) mediates the differential binding patterns of human proteins to the *S. pyogenes* strains, we applied protein affinity purification mass spectrometry (AP-MS)^16^ as schematically shown in **Figure 1B**. Based on the different *S. pyogenes* strains screened above, six M proteins (M1, M3, M5, M28, M49 and M89) were recombinantly expressed with an affinity tag. The tagged M proteins were used to affinity purify interacting plasma proteins followed by DIA-MS and filtering. Only proteins enriched log2 > 1 times (two-fold) and having an adjusted statistical *p* value of 0.05 when compared to GFP enriched proteins were considered as interactors (see **Fig. 3A** for an example **and Fig. S2)**. This filtering strategy generated a final list of 32 high confident non-redundant interactions with M1, M3, M28, M49 and M89, categorized into the same functional categories as above. As M5 had a poor protein stability and yield, it could not be used in AP-MS and was excluded from the study. The heatmap of the significant interactions in **Figure 3B** reveals five predominant column clusters and again demonstrates that the M proteins are involved in distinct protein interactions with human plasma proteins. Fibrinogen binding was prominent to M proteins of *emm* type A-C (M1 and M3), and C4BP binding to M proteins *emm* type E (M28, M49 and M89) (**Fig. 3B**) in a similar fashion as observed in the SA-MS results above and as previously suggested by Sanderson *et. al.*^13^. To detail the properties of the differential binding patterns, we constructed another correlation network plot for the 32 proteins across the five different M proteins (**Fig. 3C**). Similar to the SA-MS results, the network view shows several correlating proteins clusters (r^2^ ≥ 0.5) typically associated to the one or two of analyzed M proteins. Several of the serotype specific proteins shown above such as fibrinogen, apolipoproteins, C4BP and IgA, are strongly associated with particular M proteins. In addition, we can confirm that binding of some proteins seems to result in lower binding of other proteins (r^2^ < −0.5) such as fibrinogen-PROS and fibrinogen-C4BPα, demonstrating that the interactions captured above using SA-MS are to a large degree mediated by the M proteins. To visualize the core-interaction network between the analyzed M proteins, we selected the highly enriched protein interactions (log2 > 3 compared to GFP) to plot a schematic interaction network graph (**Fig. 3D)**. The network graph reveals that albumin, IgG1 and IgG4 are equally associated with all analyzed M proteins. Albumin is known to bind the conserved C-repeats ^14,20,23,24^ of the M protein, thus making the association of albumin with all M proteins logical. In addition, IgA2 is enriched in all M proteins although significantly more enriched to M28, which is also coupled to C4BPα, IgA1, alpha-1-antitrypsin (A1AT) and to lesser degree PROS. The cysteine residue on the C terminus of α chain of monomeric IgA has been shown to form disulfide bonds with A1AT^52^ and C4BP is known to form complex with PROS^53^. We speculate that these proteins form a larger complex mediated via human-human protein interactions on M28. In contrast, M1 and M3 typically bind fibrinogen and fibronectin, whereas M49 binds several components of the complement system, and both M49 and M89 bind PROS. In conclusion, the results from the AP-MS analysis demonstrate that M proteins play a major role in shaping the serotype-specific HP-PPI networks observed in SA-MS. Although the E type M proteins are substantially smaller compared to A-C types, their interaction networks with human plasma proteins are still surprisingly complex. As there are no structural model for any E type M interspecies protein complex, we selected M28 for further structural characterization with a particular focus on the binding with IgA and C4BP as outlined in **Figure 1B**.

**Figure 3.**
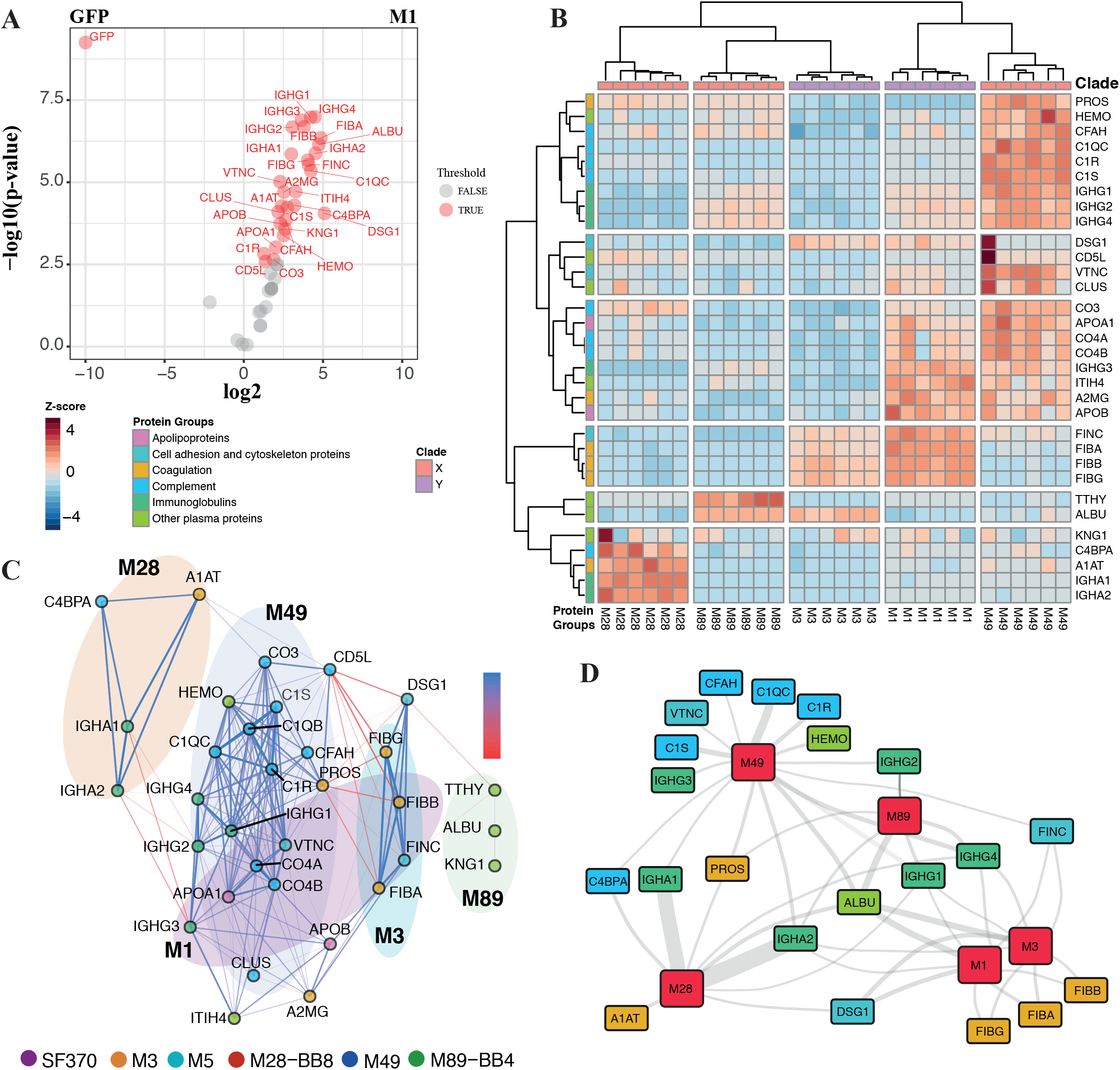
Human plasma proteins interacting with different M proteins identified by AP-MS. A) A volcano plot analysis of M1 with GFP. The DIA data was filtered against sfGFP using a log2 fold enrichment of > 1 with an adjusted P-value of 0.05 using the student t-test. The red dots represent high-confident interactors while the grey did not pass the above filtering criteria. Volcano plots for other M proteins and GFP are provided in Supplementary Figure 2 (S2). B) Cluster analysis of 32 human plasma proteins across five different M proteins for n=6 replicates. C) Person network analysis for r^2^ value of ≥ 0.5 for of 32 proteins across five M proteins. Each sphere represents a protein cluster. Blue lines represent positively correlated proteins while red lines represent mutually exclusive ones. D) Network analysis of highly significant 21 human plasma proteins across five different M proteins. These 21 proteins were log 2 > 3 enriched in M proteins as compared to GFP. The thickness of the line represents fold-change compare to GFP.

### Characterization of the M28 IgA-C4BP interaction in different local microenvironments

As we observed that IgA was significantly enriched on M28, we measured the affinity between M28 and IgA by using surface plasmon resonance (SPR). The binding of M28-IgA was compared to M1-IgA binding, which according to our observation showed very low or no IgA binding. We immobilized the M proteins (ligand) on the sensor chip and injected IgA (analyte) over them to mimic the M proteins protruding out from the bacterial surface and the immunoglobulins floating in the plasma. The kinetic analysis showed the best fit to a heterogeneous ligand model compared to a 1-1 model (**Fig. S3, A & B**). Surface heterogeneity (heterogeneous ligand) models are observed if the ligand has multiple binding sites for an analyte. Thus, an explanation for the deviations from a 1-1 fitting model could be that IgA has multiple binding sites on M28. Calculated affinity constants showed that IgA had a 3 log higher affinity for M28 as compared to M1 (K_D1_≈10^−10^ M and K_D2_≈10^−8^ M for M28 and K_D1_≈10^−7^ M and K_D2_≈10^−7^ M for M1) (**Fig. 4A-I & II**). The differences in K_D1_ and K_D2_ values of IgA towards M28 support two different binding sites on M28 for IgA, one with high and the with lower affinity. We also performed SPR analysis of the interaction with C4BP and M28 since C4BP was significantly enriched on M28 in our SA-MS and AP-MS experiments above. In this case, kinetic analysis showed a better fitting to a 1-1 model (**Fig S3, C**). Affinity constants calculated from the sensorgrams resulted in a K_D_ of 1.88 x 10^−10^, suggesting one single binding site with a high affinity between M28 and C4BP (**Fig. 4A-III**).

**Figure 4.**
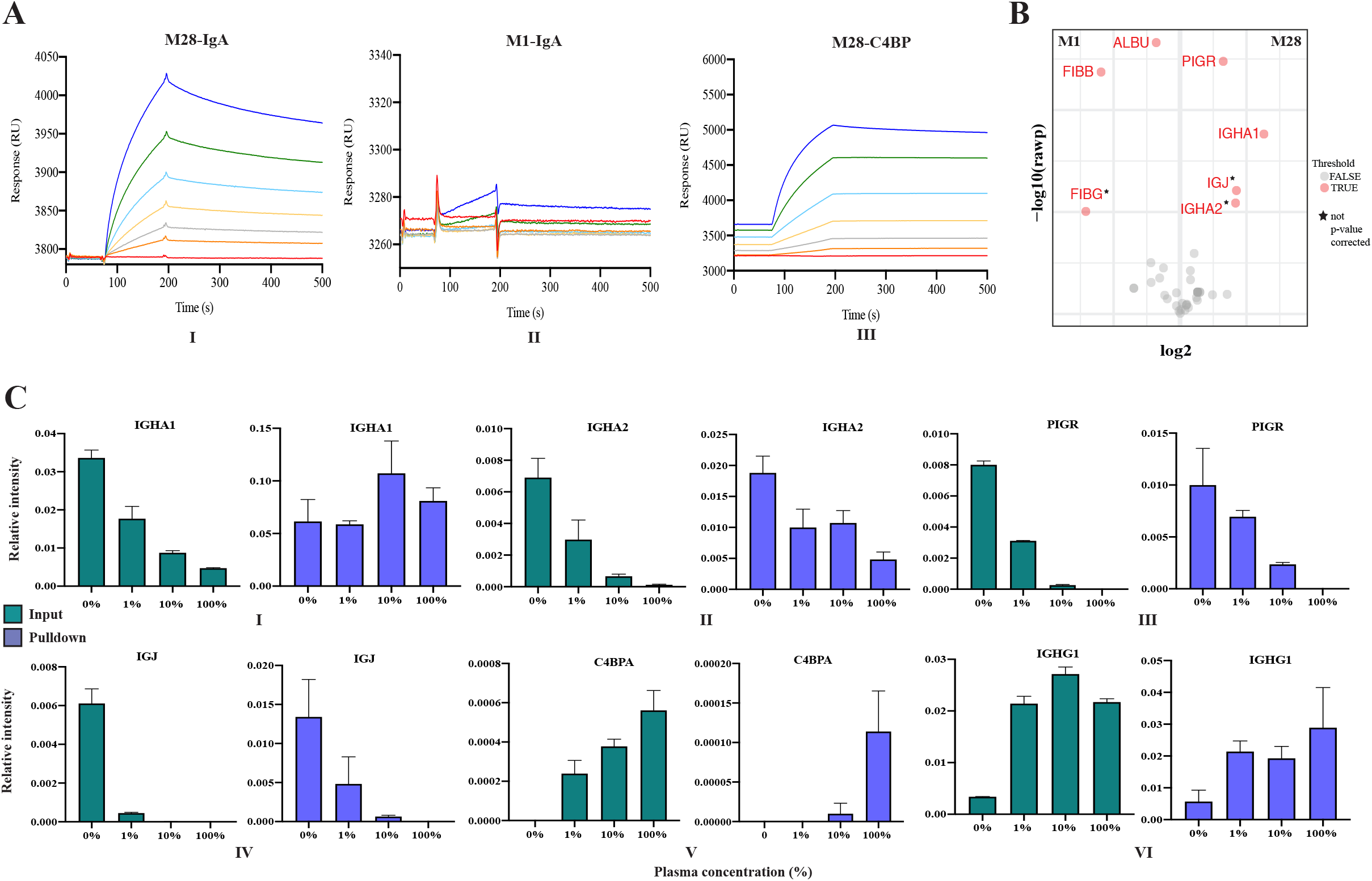
M-protein saliva-plasma interaction. A) Sensorgrams that show the response unit (RU, Y-axis) plotted as a function of time (in second, X-axis) for the interaction - (I) of M28 with IgA (K_D1_ = 3×10^−10^ and K_D2_ = 3.38×10^−8^), (II) of M1 with IgA (K_D1_ = 5.13 ×10^−7^ and K_D2_ = 3.57×10^−7^), and (III) M28 with C4BP (K_D_ of 1.88 x 10^−10^). The different color of the lines in the sensorgrams represents different concentrations of IgA and C4BP. For IgA red-0 μM, orange-0.009375, grey-0.01875, yellow-0.0375, light blue-0.075, green-0.15 and dark blue-0.3 μM. For C4BP red-0, orange-3, grey-6, yellow-12, light blue-24, green-48 and dark blue-96nM. B) A volcano plot for AP-MS of saliva with M1 and M28 to identify true human saliva proteins interacting with M28. The data was filtered using a log2 fold enrichment > 1 with an adjusted P-value of 0.05 using the student t-test. Proteins marked with stars have a non-adjusted p-values. Red dots represent high-confident interactors while grey dots did not pass the filtering criterion. C) Relative intensity of peptides (Y-axis) plotted against 0, 1, 10 and 100% plasma concentration (X-axis) mimicking vascular leakage for IGHA1 (I), IGHA2 (II), PIGR (III), IGJ (IV), C4BPA (V) and IGHG1 (VI). Green represents input samples and purple represents pulldown with M28.

Most IgA produced in the human body is secreted into the mucus membrane thereby acting as a first line defense against infections^27^. To understand how an IgA rich microenvironment alters the *S. pyogenes* M28 protein network, we quantified the protein interaction network of M1 and M28 in pooled normal human saliva by AP-MS. These experiments showed that IgA binding from saliva only occurs on M28 but not on M1 (**Fig. 4B**). Additionally, we observed co-enrichment between IgA and polymeric immunoglobulin receptor (PIGR) and IGJ (**Fig. 4B**). PIGR is known to bind polymeric IgA and IgM at the basolateral surface of epithelial cells. PIGR bound polymeric IgA undergoes transcytosis to the luminal surface where cleavage by one or more proteinases result in secretory IgA (sIgA)^52^. The J chain forms a disulfide bridge between the cysteine residues of the IgA heavy chain giving rise to multimeric IgA^52^.

Polymeric IgA is known to be prevalent in saliva while monomeric IgA and C4BP are predominantly present in plasma. *S. pyogenes* typically induce vascular leakage when localized in the upper respiratory tract^54^ thereby altering the protein composition in the host microenvironment^55^. To understand how saliva or plasma alters the *S. pyogenes* M28 protein interaction network, we quantified the protein interactions of M28 in a mixed saliva-plasma environment. These AP-MS experiments were performed using 100% saliva, 1% plasma, 10% plasma in saliva and 100% plasma to mimic conditions during a local infection followed by a systemic infection. The results show that M28 can enrich IgA1 to similar levels across all saliva or plasma mixtures, although the concentration of IgA1 is lower in plasma (**Fig 4C-I)**. In contrast, IgA2 binding to M28 predominantly occurs in saliva and decreases with the decreasing IgA2 concentration in plasma (**Fig. 4C-II**). The levels of PIGR and IGJ binding to M28 (**Fig. 4C-III & IV**) follows a similar trend, although we note proportionally higher levels of these two proteins enriched on M28 in 10% plasma compared to the input concentration (**Fig 4C-III & IV**). These results imply that M28 can bind IgA in both sIgA and monomeric form where the former is pronounced in saliva. The higher levels of AP-purified IGJ and PIGR compared to the input pool in 10% plasma environment, with nearly 18 times higher plasma protein concentration compared so saliva, suggest that the sIgA binds with high affinity which is in contrast to previously published results^56^. The mixed saliva-plasma enrichment comparison of M28 additionally revealed elevated levels of C4BP on M28 only at high plasma concentrations (**Fig. 4C-V**). Interestingly, although there were detectable levels of C4BPA in 1% plasma, there was no strong enrichment of C4BPA to M28 at this low plasma concentration. These results are surprising as the SPR analysis showed that the affinity between M28 and C4BPA was in the sub nanomolar range. Possibly this could be accounted to the fact that the levels of secretory IgA were still high at 1% plasma. The levels of IgG1 enriched on M28 seemed to increase with increase in plasma concentration (**Fig. 4C-VI**). Collectively, these results show that M28 binds secretory IgA in saliva and C4BP binding on M28 only becomes accentuated in the absence of secretory IgA.

### Structural determination of M28 with IgA and C4BP

To understand how the shorter E type M28 binds secretory IgA in saliva and monomeric IgA and C4BP in plasma, we performed targeted cross-linking mass spectrometry (TX-MS^17^) and hydrogen-deuterium mass spectrometry (HDX-MS) of M28 in complex with C4BP or the Fc-domain of IgA. As the input for TX-MS-based structural modelling, we first generated a computational model of the full-length M28, which was determined using the Rosetta comparative modeling protocol^49^ based on the previously reported model of the M1 protein^26^. This model was further used to provide protein-protein docking decoys using structures deposited in protein data bank for IgA (PDB 6LXW) and C4BP (PDB 5HYP). For TX-MS, the affinity-tag of M28 was removed, and the untagged protein was cross-linked individually in solution to either C4BP or the Fc-domain of IgA. The cross-linked peptides observed between M28 and C4BP overlapped with the interaction interface resolved using X-ray crystallography^57^ (**Fig. 5A-B, Fig. S4, ST2**). These cross-links were observed between two C4BP residues (K28 and K67 in PDB 5HYP; corresponding residues K72 and K111 in the full-length C4BPα chain) and K50 on our M28 construct (**Fig. 5A-B, ST1**). No cross-links from C4BP were observed to the crystallized M28 segment (**Fig. 5A-B**), most likely due to the lack of stereo chemical favorable lysine residues at the interaction interface. For IgA, we identified two different cross-linked sites by TX-MS, the first one was supported by four inter-protein cross-links and overlaps with the previously identified M22-based IgA-binding SAP-peptide ^27^. **(Fig. 5A, C-I, Fig. S4 & ST2)**. In addition, TX-MS also identified a novel IgA-Fc interface in the middle of M28 supported by eight high-confident inter-protein cross-links (**Fig. 5C-II, Fig. S4, ST2**). The two binding sites between IgA and M28 could result in the binding of either two single IgA-Fc’s (**Fig. 5CI-II**) or one sIgA molecule, where a dimeric IgA is bridged by a J-chain and a secretory component (**Fig. 5C-III**). Using a recently determined structure for sIgA^58^ as input for TX-MS, we showed that the binding between sIgA and M28 is supported by five unique inter-protein cross-links **(Fig. 5C-III).** The binding of sIgA onto two separate and possibly synergistic binding sites on M28 could explain why sIgA binding was more pronounced in the AP-MS experiments compared to C4BP as shown above (**Fig 4C**).

**Figure 5:**
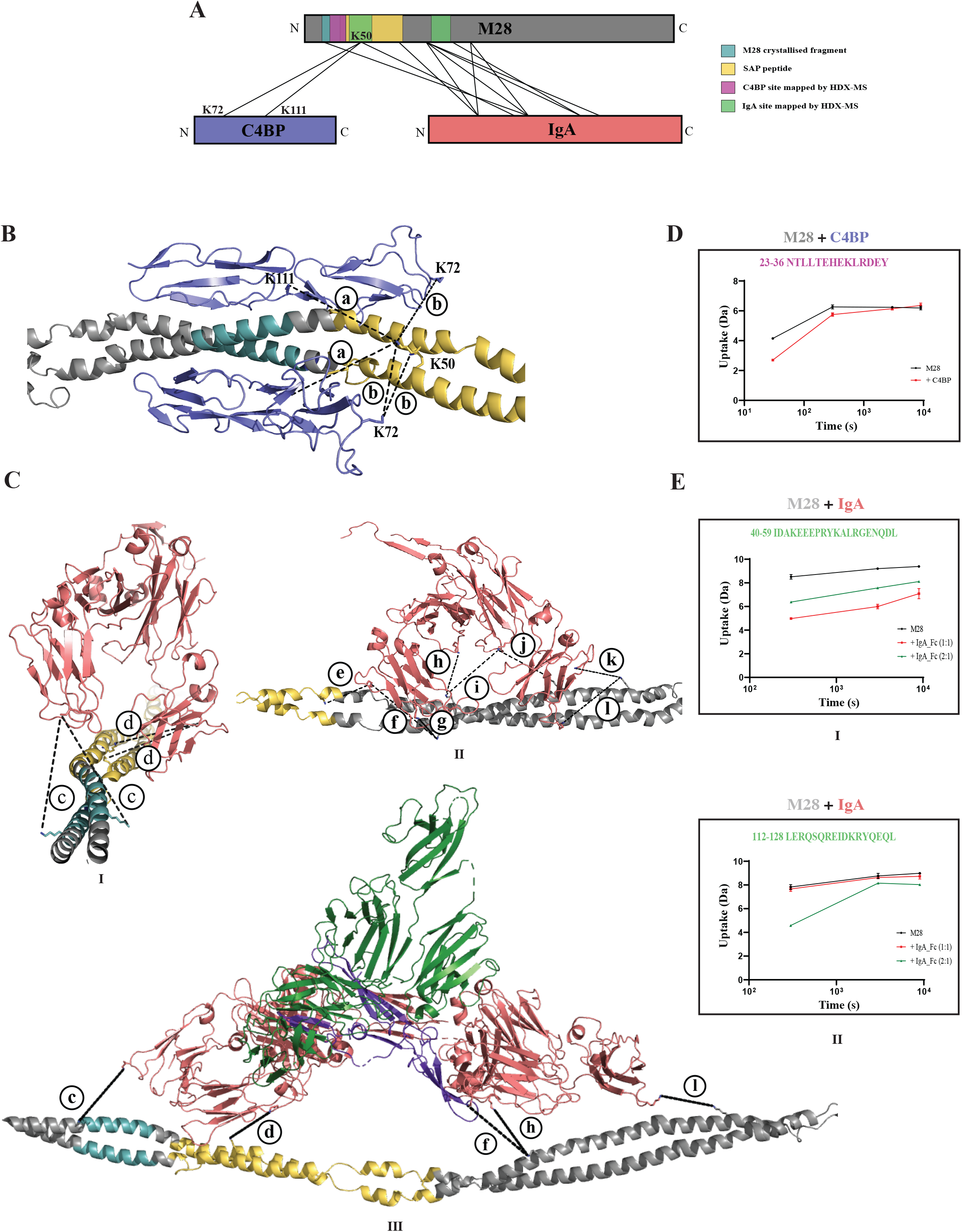
Identified interaction interfaces of C4BP and IgA on M28. A) Schematic depicting the binding regions of C4BP and IgA on M28 as identified by TX-MS and HDX-MS. B) A close-up view of the cross-linked site identified between M28 (grey helix) and C4BP (blue). The interaction interface on the crystallized M28 segment (PDB 5HYP) is shown in cyan, and the SAP-peptide interacting with the IgA Fc-domain in yellow. Cross-links are observed between lysine residues K72 and K111 (numbered based on the full-length C4BPα chain) and K50 on our M28 construct. The cross-links are depicted as dotted lines, with the labels corresponding to a given spectrum in Figure S4 and supplementary table 2 (ST2). Due to the dimeric nature of M28, several combinations of the cross-links are possible C) Close up view of the IgA-Fc binding interface on M28 identified by TX-MS. (I) The cross-linked site overlapping with the identified C4BP interaction interface and the previously identified M22-based IgA-binding SAP-peptide between the M28 (grey helix) SAP-region (yellow) and the IgA Fc-domain (red) viewed down along the helix. The cross-links are depicted as dotted lines, with the labels corresponding to a given spectrum in Figure S4 and supplementary table 2 (ST2). (II) The novel interaction site between M28 (grey helix) and the IgA Fc-domain (red). The cross-links are depicted as dotted lines, with the labels corresponding to a given spectrum in Figure S4 and supplementary table 2 (ST2). (III) The possible secretory IgA-Fc (Red) M28 model. The purple represents the J chain, and the secretory component is represented in green. The cross-links are depicted as dotted lines, with the labels corresponding to a given spectrum in Figure S4. (III) and supplementary table 2 (ST2). D) Deuterium uptake graph for the amino stretch 23-36 on M28 alone (black) and M28 and C4BP (red). E) Deuterium uptake graph for M28 and IgA – (I) 40-59 aa acid stretch on M28, the suggested high-affinity site and (II) 112-128 amino acid stretch on M28 the suggested low affinity site. Black represents M28 alone, red is M28 IgA-Fc (1:1 ratio) and green represents M28 IgA-Fc (2:1).

Complementary to the TX-MS analysis, bottom-up HDX-MS experiments were performed to track solvent protection of the interaction interface of M28 when bound to either C4BP or IgA-Fc. HDX-MS identified a 14-amino acid stretch (23-36 aa) in the HVR domain of M28 interacting with C4BP (**Fig. 5A, D)**, enclosed between the C4BP-binding site in the crystallized complex (PDB 5HYP), and the cross-linked site (K50) identified by TX-MS (**Fig. 5A**). HDX-MS analysis of the M28-IgA Fc-domain interaction showed strong protection to deuterium uptake at two distinct sites. At an M28 to IgA ratio of 1:1 the reduction in deuterium uptake was observed at the SAP-peptide and the overlapping region identified by TX-MS (**Fig. 5A, C-I, E-I**). A reduction in deuterium uptake was furthermore observed for residues 112-128 at an M28 to IgA ratio of 2:1, especially at short labeling times **(Fig. 5A, E-II)**, indicating a lower affinity site as suggested by the SPR data (**Fig 4B**). Importantly, this latter M28 site protected to deuterium uptake overlaps with the IgA-Fc interface identified by TX-MS (**Fig. 5A, C-II**). Taken together, our AP-MS data in combination with the integrative structural mass spectrometry approach, allowed us to propose two distinct models for the M28 interactions. In one model two single IgA Fc-monomers and a C4BP-moculecule would simultaneously bind to M28 (**Fig. 6A**) and in the other one the IgA-binding sites would be occupied by sIgA alone (**Fig. 6B**). The results suggest that the domain arrangement of M28 enables *S. pyogenes* to form host microenvironment dependent protein interactions. The ability to alter the protein interaction network depending on the host microenvironment allows *S pyogenes* to initiate critical immune evasion strategies in different ecological niches of relevance for both mucosal and systemic infections.

**Figure 6:**
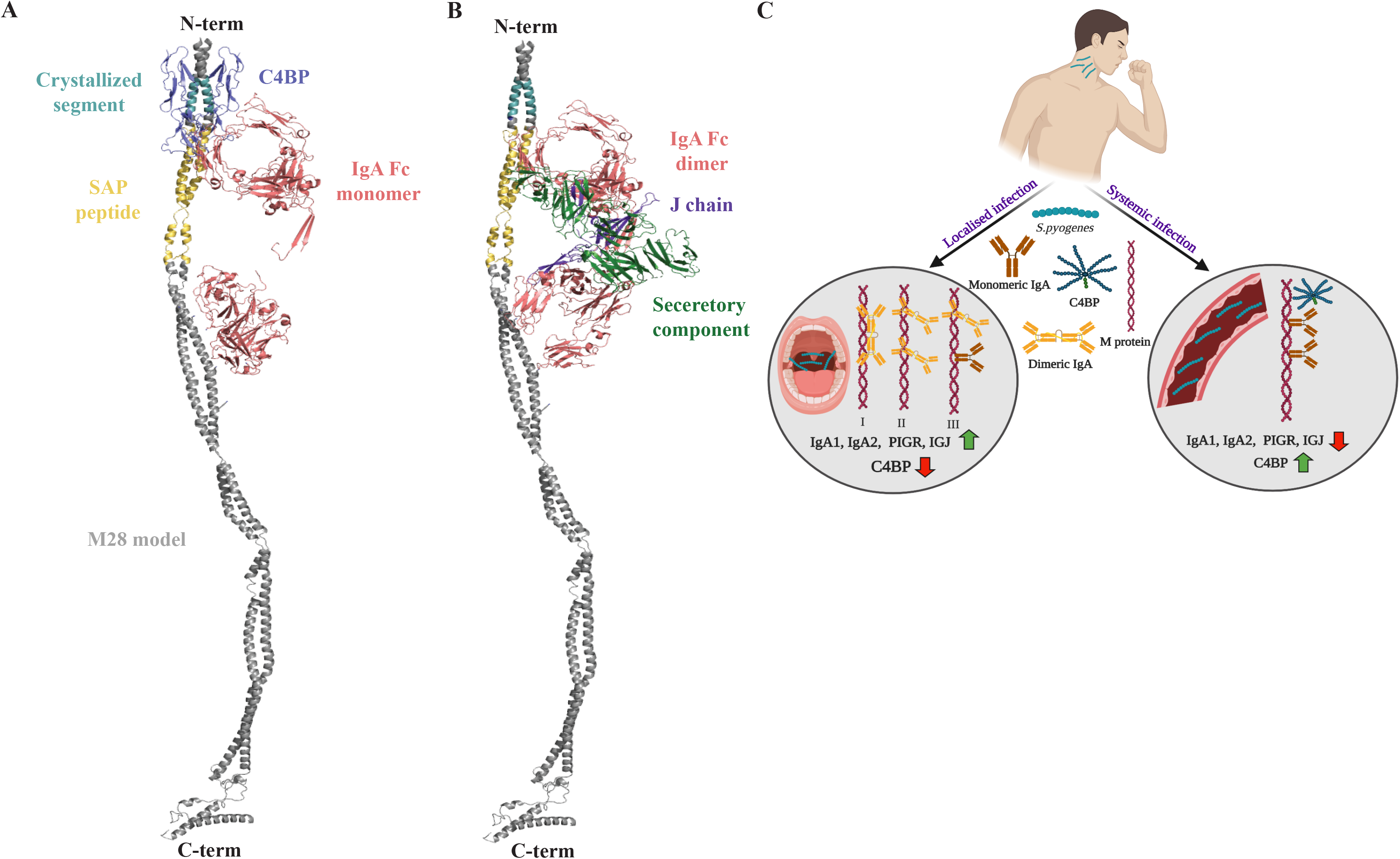
Overview of IgA-Fc and C4BP binding to M28. The homology model for the E-type M protein M28 is depicted in grey helix. Cyan represents the X-ray crystallized M28 domain while yellow is the known IgA-Fc binding SAP peptide. A) M28 model depicting the concomitant binding of C4BP (blue) and two IgA-Fc monomers (red). B) M28 model with the secretory IgA-Fc (red). Green represents the secretory component and purple represents J chain respectively. C) A schematic overview of M28 binding secretory IgA, monomeric IgA and C4BP in different microenvironment’s in case of a localized and systemic infection.

## Discussion

The clinical manifestations of *S. pyogenes* are diverse^3^. This bacterium presents itself on skin and throat causing localised infections, but can also breach the cellular layer to cause systemic infections. This forces *S. pyogenes* to adapt to different host microenvironments. In this study, we used a combination of MS-based methods to demonstrate that different streptococcal serotypes bind specifically to distinct sets of human proteins depending on serotype and local microenvironment. These interactions were in turn mediated by one of the most abundant and widely studied surface attached virulence factors, the M proteins. The newly established M-centred interaction networks recapitulated many of the previously identified M protein–human interactions and in addition highlights several so far functionally uncharacterized protein interactions. Interestingly, we note that the binding interactions were highly divergent between the analysed *emm* types. A prominent example is the binding of fibrinogen to A-C pattern and of C4BP to the E pattern M proteins. Fibrinogen is known to bind to the B repeats of the M protein^10,14,19^ and C4BP to HVR domain^32–36^ confirming that serotype-specific networks are highly dependent on the M protein domain arrangement. The binding of C4BP further strongly correlates with the binding of vitamin K-dependent protein S (PROS). As there is a possible functional redundancy between PROS and fibrinogen it cannot be excluded that E type M proteins that do not bind fibrinogen require the enrichment of an anticoagulant on their surface to evade being trapped in clots. A notion that requires further investigations. The results imply that sequence variability and the domain arrangement of M proteins can result in affinity differences to facilitate recruitment of human proteins that maximizes the chances for successful immune evasion in different microenvironments. These interactions can mediate immune evasion through different and sometimes complementary pathways. Our data suggests that once an interaction has been established with a particular protein, other plasma protein interactions are not readily formed to the same *emm* type. Possibly since some of the interacting proteins participate in similar immune evasion functions.

Both the SA-MS and AP-MS data revealed strong IgA interaction with M28, while no other serotype of M protein investigated in this study was observed to bind IgA to the same degree. The SAP peptide derived from M22^27^ was previously shown to harbor an IgA binding site. In our study, M28 is the only M protein which contains the SAP-peptide sequence **(Fig. S1).** IgA is the most abundant immunoglobulin on the mucosal surface. As *S.pyogenes* are known to localize in mucosal surfaces, hence strong binding of IgA to M28 could be warranted and likely playing a role in facilitating the bacteria to evade the first line of immune defense on the mucosal surface. In fact, the M28 serotype has been reported to be one of the leading causes of puerperal sepsis^59–62^. Persistent infections of the mucosal membrane by *S.pyogenes* can induce vascular leakage thereby providing access of the bacterium to human plasma. Here we mimic a localized infection condition followed by a systemic infection and we observe that under such circumstances, the M28 interaction network gradually changes its composition from predominant binding of secretory IgA in saliva to monomeric IgA and C4BP in plasma. This change is driven by the differences in protein concentration in the host microenvironment. However, even at higher plasma concentrations (10% plasma), secretory IgA is enriched to a higher extent to M28 compared to the input sample, whereas C4BP is not enriched to the same extent under these conditions. Typically, bacteria–host relationships are well-balanced. Sepsis is a relatively rare condition compared to uncomplicated local infections, implying that the evolution of bacteria–host relationships is predominately taking place in local host microenvironments and not in blood as previously proposed^25^. In local microenvironments, secretory IgA is the major immunoglobulin. Our results support the following three models in a mucosal niche: i) one dimeric IgA occupying both the IgA binding sites on M28; ii) two dimers binding separately to the two sites and lastly; iii) one dimer and one monomer could be engaged on M28 **(Fig. 6C)**. However, the stoichiometry of the IgA dimer binding in such a condition still remains unexplored. During the course of an infection there can be local damage of the mucosal membrane causing leakage of plasma exudate thus creating an upsurge of C4BP in the local environment^33^. Under such circumstances the bacterium is known to encounter IgG from plasma but *S. pyogenes* have many well described virulence factors like EndoS^63^, SpeB^64^, IdeS^65^ to circumvent IgG effects. This change in local microenvironment may therefore drive binding to C4BP along with monomeric IgA **(Fig. 6C)**. It has been reported that binding of both IgA and C4BP to a M protein is crucial in inhibiting phagocytosis^31^. C4BP is known to bind to the HVR of the M proteins and IgA binds a semiconservative domain adjacent to the HVR site^31^, which makes concomitant binding of C4BP and monomeric IgA to M28 plausible as previously suggested^27^. As M proteins are known to be imperfectly coiled^66^, there might also be a real possibility that the binding of one protein at a certain site might introduce conformational changes in other parts of the coiled-coil thereby affecting the affinity of proteins on another site. In this case, binding of monomeric IgA might induce a conformational change that promotes binding of C4BP to the HVR of M28, in a manner similar to what has been shown for increased C4BP-binding to the streptococcal surface mediated by IgG^67,68^. This could also explain the strong coupling seen between IgA and C4BP in the SA-MS and AP-MS experiments. We propose that M28 binds either secretory IgA or monomeric IgA and C4BP depending whether they cause a localized infection or a systemic infection **(Fig. 6C)**. The structural model presented here is consistent with our finding of M28’s dimeric IgA binding and concomitant binding to monomeric IgA and C4BP but in different ecological niches.

## Acknowledgements

Gene cloning, protein expression, and purification of the M1 protein, sfGFP was performed at the Lund Protein Production Platform (LP3), Lund University, Sweden (www.lu.se/lp3). Daniel Hatlem from Dirk Linke’s lab is highly acknowledged for his guidance during recombinant protein purification. We thank Oonagh Shannon for access to the different *S.pyogenes* serotypes. Support from the Swedish National Infrastructure for Biological Mass Spectrometry (BioMS) is gratefully acknowledged. The illustration 1B and 6C was created using BioRender.com and 5A was prepared in xVis.

This research was supported by the Viral and Bacterial Adhesin Network Training (ViBrANT) Program funded by the European Union’s HORIZON 2020 Research and Innovation Program under the Marie Sklodowska-Curie Grant Agreement No 765042 to JM and DL, the Swedish Research Council (2019-01646) to JM, the Foundation of Knut and Alice Wallenberg (2016.0023 and 2019.0353) to JM and LM and Österlunds Stiftelse to JM and RL. HK was supported by the Swiss National Science Foundation (early postdoc mobility grant no. P2ZHP3_191289).

---

**Supplementary figure 1**. **Multiple sequence alignment of mature M proteins**. Multiple sequence alignment of the amino acid sequence of M1, M3, M28, M49 and M89 proteins recombinantly expressed and used for APMS experiments. The yellow highlighted region represents the SAP peptide sequence on M28. The red region represents the novel identified IgA-Fc binding site on M28.

**Supplementary figure 2**. **Volcano plot for APMS of M proteins with human plasma**. Volcano plot analysis of different M proteins with GFP with a filtering criterion of log2 fold enrichment > 1 with an adjusted P value of 0.05 using the student t-test. A) GFP and M3, B) GFP and M28, C) GFP and M49 and D) GFP and M89.

**Supplementary figure 3**. **Kinetic analysis of IgA binding to immobilized M28 and M1 and C4BP binding to M28 fitted to different models.** A) IgA binding for immobilized M28 fitted to (I) 1-1 model and (II) heterogeneous ligand model. B) IgA binding for immobilized M1 fitted to (I) 1-1 model and (II) heterogeneous ligand model. C) C4BP binding for immobilized M28 fitted to (I) 1-1 model and (II) heterogeneous ligand model.

**Supplementary figure 4. Individual spectra for identified cross-linked peptides.** Spectra A-B) are for the M28 – C4BP interface, spectra C-D) M28 – IgA interface overlapping with the identified C4BP interaction interface and the previously identified M22-based IgA-binding SAP-peptide and spectra E-L) for the novel M28 – IgA interface. The cross-linked peptides are indicated at the top of each spectrum. The fragments indicated in red and blue are from the individual parent peptides, whereas the fragments in green arise from a fragmented cross-linked peptide. The intensity for each spectrum is shown on the y-axis and the m/z ratio on the x-axis.

**Supplementary table 1 (ST1)**. **M protein sequences and UniProt IDs**. Protein sequences of different M proteins expressed recombinantly along with their UniProt IDs are represented in the table. The initiating methionine residue is marked in red and the colored sequence represents the tag incorporated in the protein.

**Supplementary table 2 (ST2)**. **Cross-linked peptide list**. List of peptides from M28 cross-linked to peptides from either C4BP or IgA-Fc. The cross-linked lysine residues (K) are bolded.

